# OPUS-Fold3: a gradient-based protein all-atom folding and docking framework on TensorFlow

**DOI:** 10.1101/2022.08.31.506128

**Authors:** Gang Xu, Zhenwei Luo, Ruhong Zhou, Qinghua Wang, Jianpeng Ma

## Abstract

For refining and designing protein structures, it is essential to have an efficient protein folding and docking framework that generates a protein 3D structure based on given constraints. In this study, we introduce OPUS-Fold3 as a gradient-based, all-atom protein folding and docking framework, which accurately generates 3D protein structures in compliance with specified constraints, such as a potential function as long as it can be expressed as a function of positions of heavy atoms. Our tests show that, for example, OPUS-Fold3 achieves performance comparable to pyRosetta in backbone folding, and significantly better in side-chain modeling. Developed using Python and TensorFlow 2.4, OPUS-Fold3 is user-friendly for any source-code level modifications and can be seamlessly combined with other deep learning models, thus facilitating collaboration between the biology and AI communities. The source code of OPUS-Fold3 can be downloaded from http://github.com/OPUS-MaLab/opus_fold3. It is freely available for academic usage.

## Introduction

Generating a protein 3D structure is a crucial task in protein structure modeling. Unlike some structure prediction methods, such as AlphaFold2 [1], which produce a specific structure based on multiple sequence alignment (MSA), a protein folding framework that can incorporate multiple sources of constraints, such as experimental and physicochemical information, local environment-derived constraints, and constraints obtained through other methods, is useful in many scenarios of application.

Over the past few decades, numerous folding frameworks have been proposed to address this issue [2-5]. For example, to generate an all-atom model using contact-map-based methods, such as RaptorX-Contact [6], one could utilize Crystallography and NMR System (CNS) [2]. Other methods like CONFOLD [7] and pyconsFold [8] also employ CNS suite [2] as their underlying folding frameworks. Recently, trRosetta [9] proposed a set of constraints that include both distance and orientation distribution information to generate 3D structures based on the pyRosetta folding framework [3, 4].

In our previous studies, we developed a protein folding framework, named OPUS-Fold2, to model the backbone [10] and side chains [11] separately using distance and orientation distribution constraints proposed by trRosetta [9]. However, folding the backbone exclusively [10] and modeling the side chains with a fixed backbone afterward [11] have limited applications. Here, in OPUS-Fold3, we propose an all-atom folding framework that enables simultaneous adjustment of the backbone and side chains during the folding process, which is beneficial for tasks requiring all-atom refinement for example. Moreover, OPUS-Fold3 can handle constraints between the receptors and ligands, making it suitable as a protein docking framework.

Moreover, recently, Giri *et al*. [12] demonstrated that reconstructing protein structures from cryo-EM density maps could benefit from integrating with the latest protein structure prediction techniques. Traditional methods, such as EM-Fold [13] and Phenix [14], implement optimization algorithms for fitting the protein structures into experimental cryo-EM density maps. These programs are largely implemented by homegrown libraries. However, current state-of-the-art structure prediction algorithms are mostly implemented by deep learning frameworks such as Tensorflow [15] and Pytorch [16], with which the traditional methods are hard to integrate. We hence implement a new fitting routine in our Tensorflow-based folding framework OPUS-Fold3, in which experimental density maps can be included as a differentiable constraint and be integrated with other constraints such as those from structure prediction methods to jointly guide the folding process.

## Methods

### Datasets

To assess the performance on monomer target, we employ the CAMEO60 [17] dataset, which comprises 60 monomer hard targets released between January and July 2020 on the CAMEO website [18]. For oligomer target evaluation, we compile a CAMEO75o dataset containing 75 targets (each with two peptide chains and fewer than 1000 residues in length) sourced from CAMEO-Homo, CAMEO-Hetero, and CAMEO93o [19]. In the PDB files, the first peptide chain is designated as the “receptor,” while the last peptide chain is identified as the “ligand.” To evaluate OPUS-Fold3’s performance as a framework for utilizing experimental density maps, we gather eight single-particle targets and their corresponding density maps, with resolutions ranging from 2.6 to 3.4 □.

### OPUS-Fold3

OPUS-Fold3 is a gradient-based, all-atom folding and docking framework. Folding-related variables in OPUS-Fold3 encompass backbone torsion angles (*Φ, Ψ* and *Ω*) and side-chain dihedral angles (*X*_*1*,_ *X*_*2*,_ *X*_*3*,_ and *X*_*4*_) for all residues.

Docking-related variables in OPUS-Fold3 consist of six parameters in the rotation matrix and three parameters in the translation matrix. In this context, we employ the distance and orientation distribution constraints proposed by trRosetta [9]. Additional constraints can be readily integrated into OPUS-Fold3, as long as they can be expressed as functions of heavy atoms’ positions.

The loss function of trRosetta-style constraints is defined as follows:

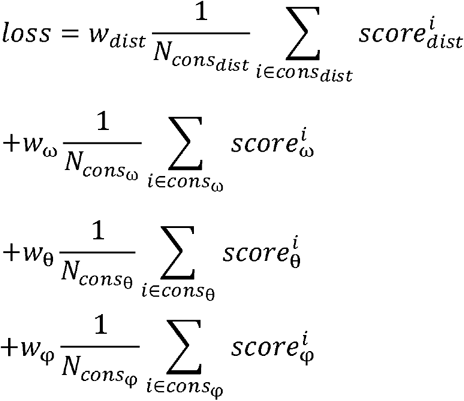

In accordance with the definitions in trRosetta [9], *cons*_*dist*_ represents the set of distance constraints (C_β_-C_β_ distance), where P_4≤dist <20_ ≥0.05.*cons*_ω_and *cons*_*θ*_are the collections of ω (C_α1_-C_β1_-C_β2_-C_α2_) and θ (N_1_-C_α1_-C_β1_-C_β2_ and N_2_-C_α2_-C_β2_-C_β1_) constraints, respectively, with *P*_*contact*_ ≥0.05. *cons*_φ_ denotes the collection of φ (C_α1_-C_β1_-C_β2_ and C_α2_-C_β2_-C_β1_) constraints, where *P*_*contact*_ ≥ 0.65.The weights for each term, *W*_*dist*_, *W*_ω_, *W*_θ_ and *W*_φ_, are set at 10, 8, 8, and 8, respectively. The distance and orientation distributions are converted to the energy terms by the following equations:

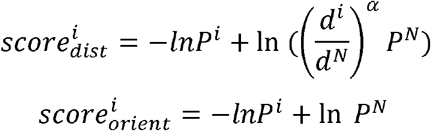

The α value is set at 1.57 [20]. Similar to trRosetta [9], the reference state for the distance distribution is the probability of the *N*th bin [19.5, 20], while the reference state for the orientation distribution is the probability of the last bin [165°, 180°]. In this context, *P*^*i*^ denotes the probability of the *i*th bin, and *d*^*i*^ represents the distance of the *i*th bin. Cubic spline curves are generated to make the terms differentiable.

Additionally, the Ramachandran scoring term [21] is employed to regulate the backbone torsion angles (*Φ* and Ψ). A radial basis function is utilized to render the probabilities differentiable. The Omega scoring term is applied for regulating the backbone torsion angle (Ω). The weights for the Ramachandran scoring term and the Omega scoring term are set at 0.1 and 0.05, respectively.

In OPUS-Fold3, the backbone modeling incorporates the original trRosetta-style constraints from trRosetta [9], the Ramachandran scoring term, and the Omega scoring term into the loss function. For side-chain modeling, the modified trRosetta-style constraints proposed by OPUS-Rota4 [11] are introduced. Specifically, four sets of constraints are included for each side-chain dihedral angle (*X*_*1*,_ *X*_*2*,_ *X*_*3*,_ and *X*_*4*_), with the corresponding side-chain atoms needed to calculate the dihedral angle defined as pseudo-C_α_ and C_β_. The detailed pseudo-C_α_ and C_β_ for each side-chain dihedral angle can be found in Supplementary Table S1.

Within a peptide chain, the backbone of each residue is sequentially generated based on the backbone torsion angles (*Φ*, Ψ and Ω), followed by the construction of side chains using the side-chain dihedral angles (*X*_*1*,_ *X*_*2*,_ *X*_*3*,_ and *X*_*4*_). During the docking process, a rotation matrix containing six parameters and a translation matrix containing three parameters are employed to determine the relative positions between the receptor and ligand. Furthermore, the coordinates of all atoms in the ligand are transformed using the aforementioned matrices.

OPUS-Fold3 is built on TensorFlow 2.4 [15], and the loss function is optimized using the Adam [22] optimizer with an initial learning rate of 0.5.

### Density Map from Atomic Structure

We employ a Gaussian mixture model to generate an electron density map from a given atomic structure. To simplify computation, the electron density of each atom is modeled by a single Gaussian with a standard deviation σ. For a protein structure with N atoms, its electron density map can be expressed as

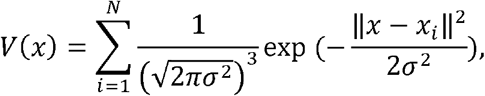

where *x* ∈ ℝ^3^ is a grid point in 3D space, and *x*_*i*_ ∈ ℝ^3^ is the coordinate of

*i*th atom in the protein structure, 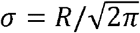 with *R* being the resolution of simutlated map is the standard deviation of the gaussian. The computation of the electron density map is differentiable with respect to the 3D atomic coordinates and can be efficiently implemented using the *tensor_scatter_nd_add* function in TensorFlow. As the Gaussian function diminishes rapidly over space, we first sample the Gaussian of each atom within a small 7x7x7 cube and then add the sampled Gaussian of each atom to the corresponding location in a large 3D array that can store the entire 3D structure.

The computational complexity of this process scales linearly with respect to the number of atoms and can be easily parallelized. The fit between the atomic structure and its experimental density map can then be achieved by maximizing the correlation between the simulated density map and the experimental density map. Let the experimental density map be *V*_o_(*x*),and the coordinates of the atomic structure be *x*_1_,…,*x*_N_; the fitting loss function can be expressed as

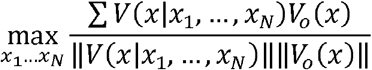

where ‖*V*(*x*|*x*_1_,…,*x*_*N*_)‖is the norm of the simulated density map, and ‖*V*_*o*_(*x*) ‖ is the norm of the experimental density map.

### Performance Metrics

We employ TM-score [23] to evaluate the accuracy of the predicted backbone. The mean absolute error (MAE) of *X*_*1*,_ *X*_*2*,_ *X*_*3*,_ and *X*_*4*_ is used to assess the accuracy of the predicted side chains. Additionally, ACC serves as a representation of the percentage of correct predictions with a tolerance criterion of 20° for all side-chain dihedral angles (from *X*_*1*_ to *X*_*4*_). ModelZ [24] is utilized to measure the fitting of the atomic structure and the experimental density map. Side-chain Z-scores are considered good when they are positive, with higher values indicating better matching between the side-chain conformation at its placed location and the map compared to slightly displaced locations [24]. Indicators (CCMapPDB, CCmain, and CCside) in Phenix [14] are also employed to measure the correlation between the atomic structure and the experimental density map.

## Results

### Performance on Backbone Folding

In Table 1, we compare the backbone folding performance of OPUS-Fold3 to the pyRosetta folding protocol in trRosetta [9] on the CAMEO60 dataset, using identical constraints as inputs. The results indicate that OPUS-Fold3 achieves performance comparable to pyRosetta, whether using predicted constraints from OPUS-Contact [10] or actual constraints derived from the corresponding PDB file. In this case, OPUS-Fold3 utilizes predicted backbone torsion angles (*Φ* and *Ψ*) from OPUS-TASS2 [10] as its initial state. When employing random values as the initial backbone torsion angles (*Φ* and *Ψ*), performance is slightly reduced (OPUS-Fold3 (random) in Table 1).

**Table 1.** The TM-score of each method on CAMEO60. “Predicted” refers to the results obtained using distance and orientation constraints predicted by OPUS-Contact. “PDB” indicates the results using the actual distance and orientation constraints derived from the corresponding PDB file. Best results for each experiment are shown in boldface.

|  | pyRosetta | OPUS-Fold3 | OPUS-Fold3 (random) |
| --- | --- | --- | --- |
| Predicted | **0.618** | 0.612 | 0.606 |
| PDB | 0.983 | **0.987** | 0.954 |

We provide examples of the backbone folding processes for OPUS-Fold3 and OPUS-Fold3 (random), using real constraints derived from the corresponding PDB file on monomer target 2020-01-04_00000019_1, in Supplementary Figure S1. The results demonstrate that when using predicted backbone torsion angles (*Φ* and *Ψ*) as the initial state, OPUS-Fold3 achieves better outcomes with fewer optimization epochs compared to OPUS-Fold3 (random). For further illustration, we display some intermediate structures during the backbone folding process of OPUS-Fold3 in Figure 1, and the folding trajectory of OPUS-Fold3 (random) is presented as a movie in Supplementary Video S1.

**Figure 1.**
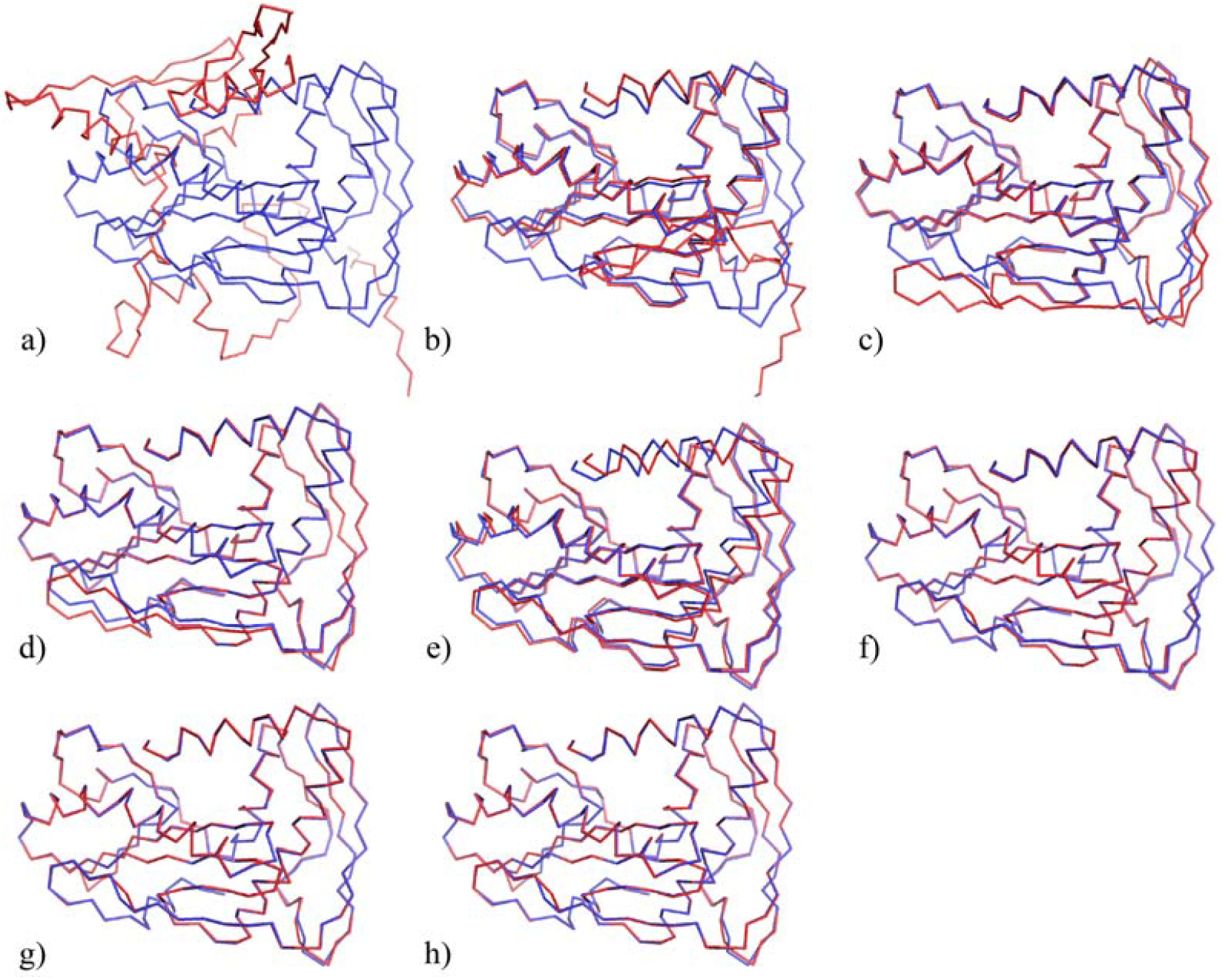
The intermediate structures of target 2020-01-04_00000019_1 (with a length of 226 residues) during the backbone folding process of OPUS-Fold3. The red structures represent the predicted intermediate structures, while the blue structure corresponds to the native state. Images a)-g) display the predicted intermediate structures at epochs 0, 200, 400, 600, 1200, 1800, and 2400, respectively. Image h) showcases the final prediction.

In Table 2, we compare the accuracy of side chain predictions from pyRosetta and OPUS-Fold3, where the side chains of OPUS-Fold3 are constructed using our side-chain modeling method, OPUS-Mut [25]. The results reveal that OPUS-Fold3’s predicted side chains significantly outperform those from pyRosetta in terms of MAE (*X*_*1*_), MAE (*X*_*2*_), MAE (*X*_*3*_), MAE (*X*_*4*_). For instance, the percentage of correct side-chain dihedral angle predictions with a 20° tolerance criterion (ACC) for pyRosetta is 42.94%, while it is 52.32% for OPUS-Fold3.

**Table 2.**
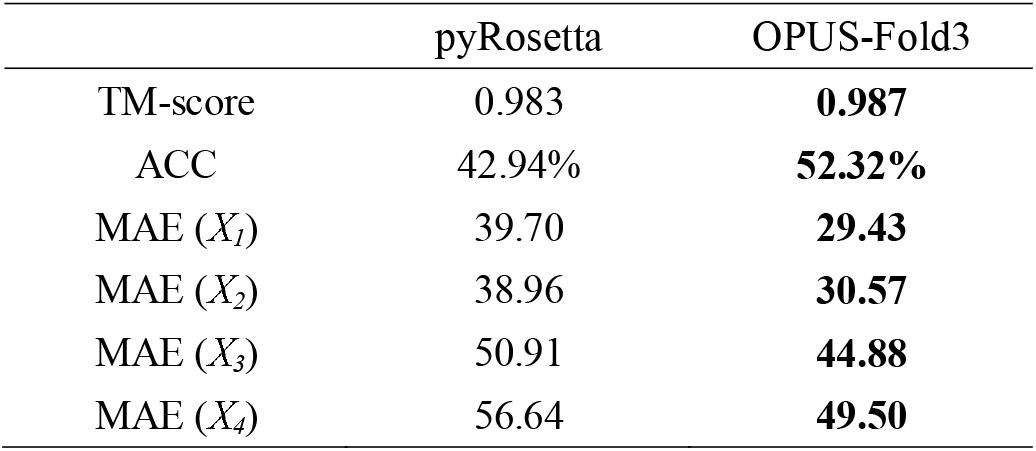
The accuracy of the predicted side chains for both pyRosetta and OPUS-Fold3. The backbones are obtained using the real constraints derived from the corresponding PDB file. Best results for each experiment are shown in boldface.

### Performance on Side-chain and All-atom Folding

In Table 3, we present the side-chain and all-atom modeling results of OPUS-Fold3 on the CAMEO60 dataset. We use real constraints derived from the corresponding PDB files to assess the performance of OPUS-Fold3 by examining the differences between the predicted structures and their native counterparts. Random values are employed as the initial side-chain dihedral angles (*X*_*1*,_ *X*_*2*,_ *X*_*3*,_ and *X*_*4*_).

**Table 3.**
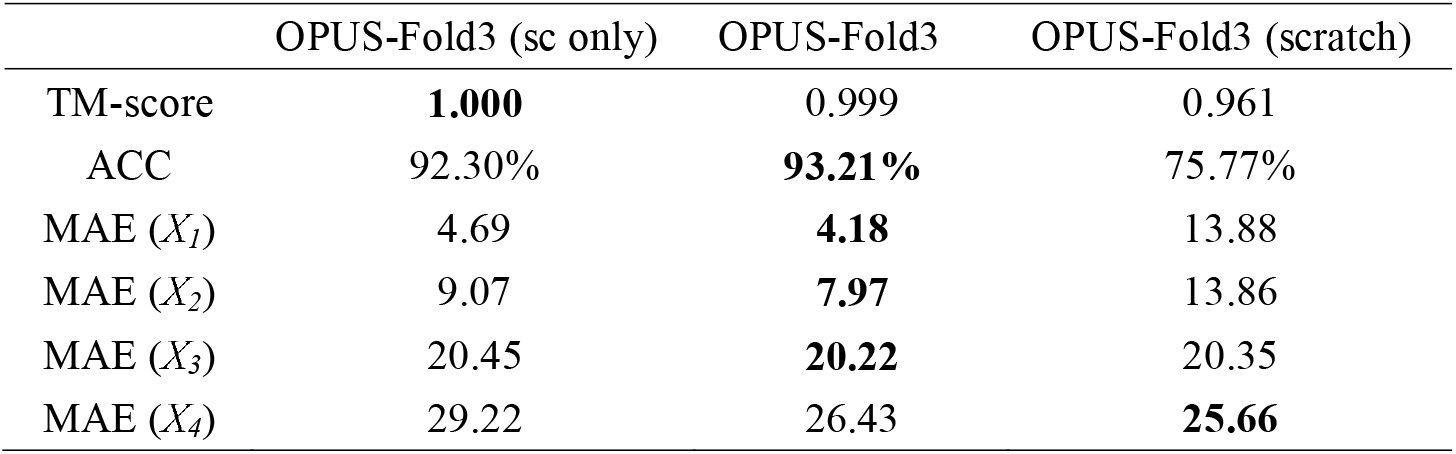
The side-chain and all-atom modeling results of OPUS-Fold3 on CAMEO60. “OPUS-Fold3 (sc only)” refers to the procedure that models the side chains with a fixed native backbone. “OPUS-Fold3” denotes the procedure that models the side chains with a fixed native backbone for the first 2400 epochs and relaxes all atoms for the final 600 epochs. “OPUS-Fold3 (scratch)” represents the procedure that models the backbone and side chains simultaneously from scratch. For each procedure, the real distance and orientation constraints derived from the corresponding PDB file are used. In OPUS-Fold3 (scratch), random values are set as the initial backbone torsion angles (*Φ* and *Ψ*). Best results for each experiment are shown in boldface.

When modeling side chains with a fixed native backbone (OPUS-Fold3 (sc only) in Table 3), the results demonstrate that the predicted side chains are highly similar to their native counterparts. This indicates the effectiveness of OPUS-Fold3 in delivering the corresponding side-chain conformation for given constraints. As an example, the folding trajectory of OPUS-Fold3 (sc only) on monomer target2020-02-15_00000234_1 (with a length of 338 residues) is showcased as a movie in Supplementary Video S2.

The results also indicate that when modeling side chains with a fixed native backbone for the first 2400 epochs, and allowing all atoms to relax during the last 600 epochs (OPUS-Fold3 in Table 3), the performance of side-chain modeling improves. This suggests that backbone relaxation may be beneficial for side-chain reconstruction.

When simultaneously modeling the backbone and side chains from scratch (OPUS-Fold3 (scratch) in Table 3), the backbone folding performance surpasses that of modeling the backbone exclusively (OPUS-Fold3 (random) in Table 1). However, the side-chain modeling accuracy declines, signifying that the correct backbone conformation is crucial for side-chain modeling.

### Increasing the Correlation between the Side-chain Atomic Structure and Density Map

To evaluate the performance of OPUS-Fold3 as an underlying framework for utilizing experimentally determined density maps, we fix the native backbone from the PDB file and refine the side chains according to the density map. Meanwhile, we only adopt the fitting loss derived from the experimental density map as our loss function. As displayed in Table 4, after refinement, the side chain Z-scores calculated by ModelZ become more positive, indicating a better match between the side-chain conformation and the map at its current location. Moreover, most of the indicators (CCMapPDB, CCmain, and CCside) calculated by Phenix also demonstrate improved correlation between the atomic structure and density map after refinement. In Figure 2, we present some examples of OPUS-Fold3 refinement results. The side chains of OPUS-Fold3 refined structures fit better into the densities, confirming that the OPUS-Fold3 refinement results enhance the correlations between experimental density maps and side chains, which means OPUS-Fold3 is capable of utilizing experimental cryo-EM maps as one of the sources of constraints for structure folding. Combined with other constraints, it may form a brige between cryo-EM structural determination and computational structure prediction techniques.

**Table 4.** The correlation between the side-chain atomic structures and their corresponding experimental density maps. The results at the top are calculated using the original atomic structure in each PDB file, while the results (“_fold3”) at the bottom are calculated using the atomic structure after refinement. Best results for each experiment are shown in boldface.

|  | resolution<br>n (Å) | # chains | #<br>residues | ModelZ | CCMapPD<br>B | CCmain | CCside |
| --- | --- | --- | --- | --- | --- | --- | --- |
| 7uwf | 2.7 | 4 | 1750 | 2.63 | 0.76 | 0.83 | 0.80 |
| 7uwf_fold3 |  |  |  | **2.80** | 0.76 | 0.83 | 0.80 |
| 7wqx | 2.7 | 1 | 501 | 1.40 | 0.41 | 0.65 | 0.62 |
| 7wqx_fold3 |  |  |  | **2.00** | **0.44** | **0.68** | **0.67** |
| 7yim | 2.6 | 1 | 592 | 2.05 | 0.59 | 0.69 | 0.66 |
| 7yim_fold3 |  |  |  | **2.36** | **0.60** | **0.70** | **0.68** |
| 8a3b | 3.1 | 7 | 1792 | 1.85 | 0.57 | 0.73 | 0.71 |
| 8a3b_fold3 |  |  |  | **2.20** | **0.59** | 0.73 | **0.73** |
| 8dm8 | 2.68 | 2 | 794 | 2.29 | 0.75 | 0.86 | 0.82 |
| 8dm8_fold3 |  |  |  | **2.53** | **0.76** | 0.86 | **0.83** |
| 8epa | 3.4 | 5 | 637 | 1.74 | 0.68 | 0.76 | 0.75 |
| 8epa_fold3 |  |  |  | **2.08** | **0.72** | **0.77** | **0.77** |
| 8eqj | 3 | 2 | 384 | 2.09 | 0.77 | 0.70 | 0.64 |
| 8eqj_fold3 |  |  |  | **2.60** | **0.79** | **0.71** | **0.66** |
| 8hdo | 2.87 | 5 | 1028 | 2.09 | 0.63 | 0.77 | 0.73 |
| 8hdo_fold3 |  |  |  | **2.52** | **0.66** | **0.78** | **0.76** |

**Figure 2.**
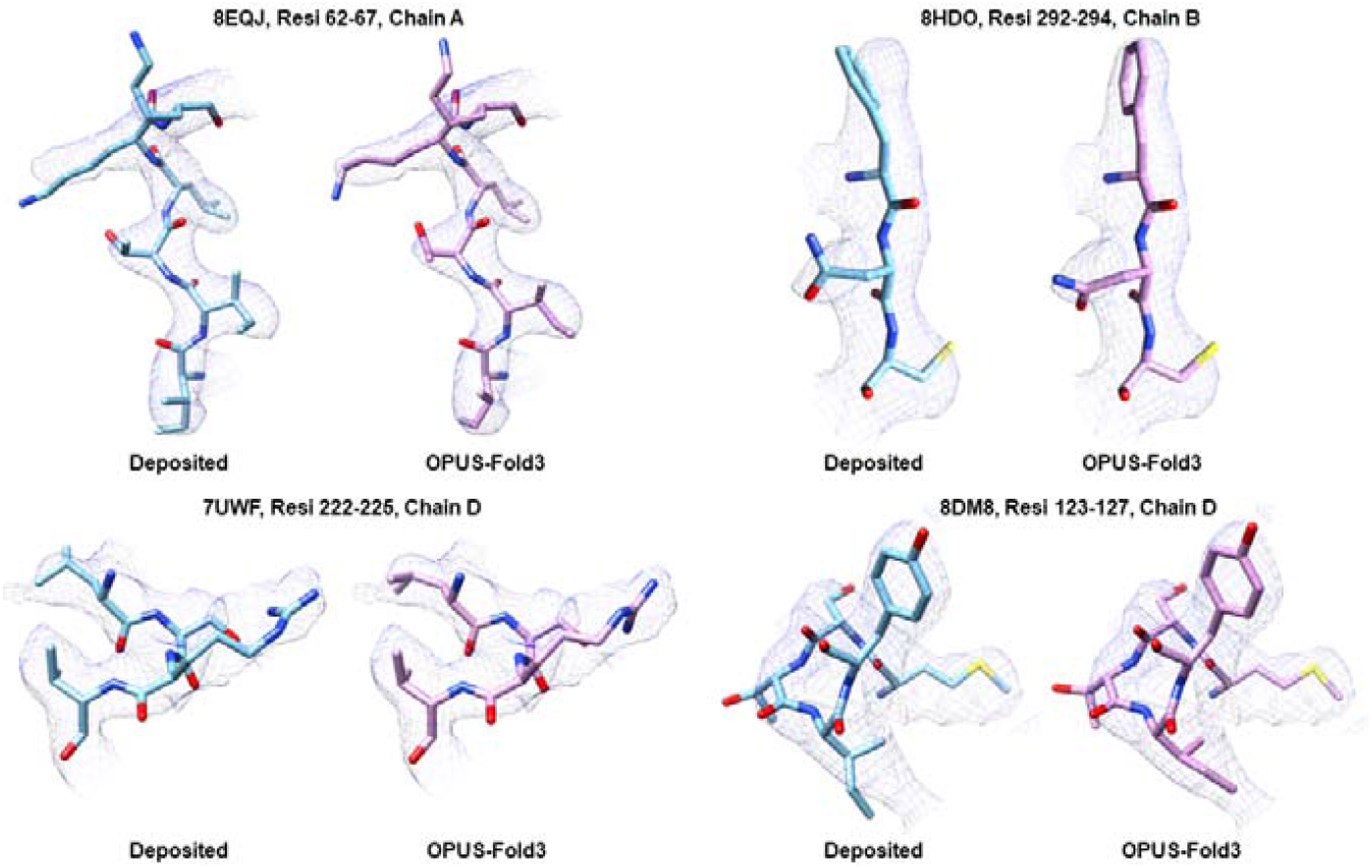
Comparison between density maps and the side chains of the deposited structures and the structures refined by OPUS-Fold. a) The structures are from residues 62-67 of chain A with PDB code 8EQJ. b) The structures are from residues 292-294 of chain B with PDB code 8HDO. c) The structures are from residues 222-225 of chain D with PDB code 7UWF. d) The structures are from residues 123-127 of chain D with PDB code 8DM8. The deposited structures are colored in blue, while the structures refined by OPUS-Fold3 are colored in violet.

In current version of OPUS-Fold3, the Rotamer outliers of the model after refinement sometimes become higher. For example, the Rotamer outliers of 7uwf is 0%, and that of 7uwf_fold3 is 21.69%. This may be caused by the inaccurate guidance from the density map alone at relatively low resolution regions. Therefore, in future work, more terms regarding geometry restraints will be included in the loss function for better structural refinement. However the current results still show that OPUS-Fold3 successfully maximizes the model-map correlation following the fitting loss term, which demonstrates the possibility of OPUS-Fold3 as an optimization framework for utilizing the information from density map.

### Performance on Protein-protein Docking

In Table 5, we present the protein-protein docking results of OPUS-Fold3 on CAMEO75o using the real constraints derived from the corresponding PDB-file. The results demonstrate that OPUS-Fold3 can deliver the correct docking pose when the backbones of the receptor and ligand are known (OPUS-Fold3 in Table 5). We display some intermediate structures during the protein-protein docking process of OPUS-Fold3 on hetero-oligomer target 7SPP in Figure 3.

**Table 5.** The protein-protein docking results of OPUS-Fold3 on CAMEO75o. “OPUS-Fold3” denotes the procedure that models the docking pose with fixed native backbones of receptor and ligand. “OPUS-Fold3 (bbfold)” represents the procedure that simultaneously models the backbones of receptor and ligand, and the docking pose between them from scratch. “OPUS-Fold3 (aafold)” refers to the procedure that simultaneously models all atoms of receptor and ligand, and the docking pose between them from scratch. For each procedure, the real distance and orientation constraints derived from the corresponding PDB file are used. In OPUS-Fold3 (aafold), random values are set as the initial backbone torsion angles (*Φ* and *Ψ*) and side-chain dihedral angles (*Χ*_*1*_, *Χ*_*2*_, *Χ*_*3*_, and *Χ*_*4*_).

|  | TM-score |
| --- | --- |
| OPUS-Fold3 | 0.994 |
| OPUS-Fold3 (bbfold) | 0.918 |
| OPUS-Fold3 (aafold) | 0.933 |

**Figure 3.**
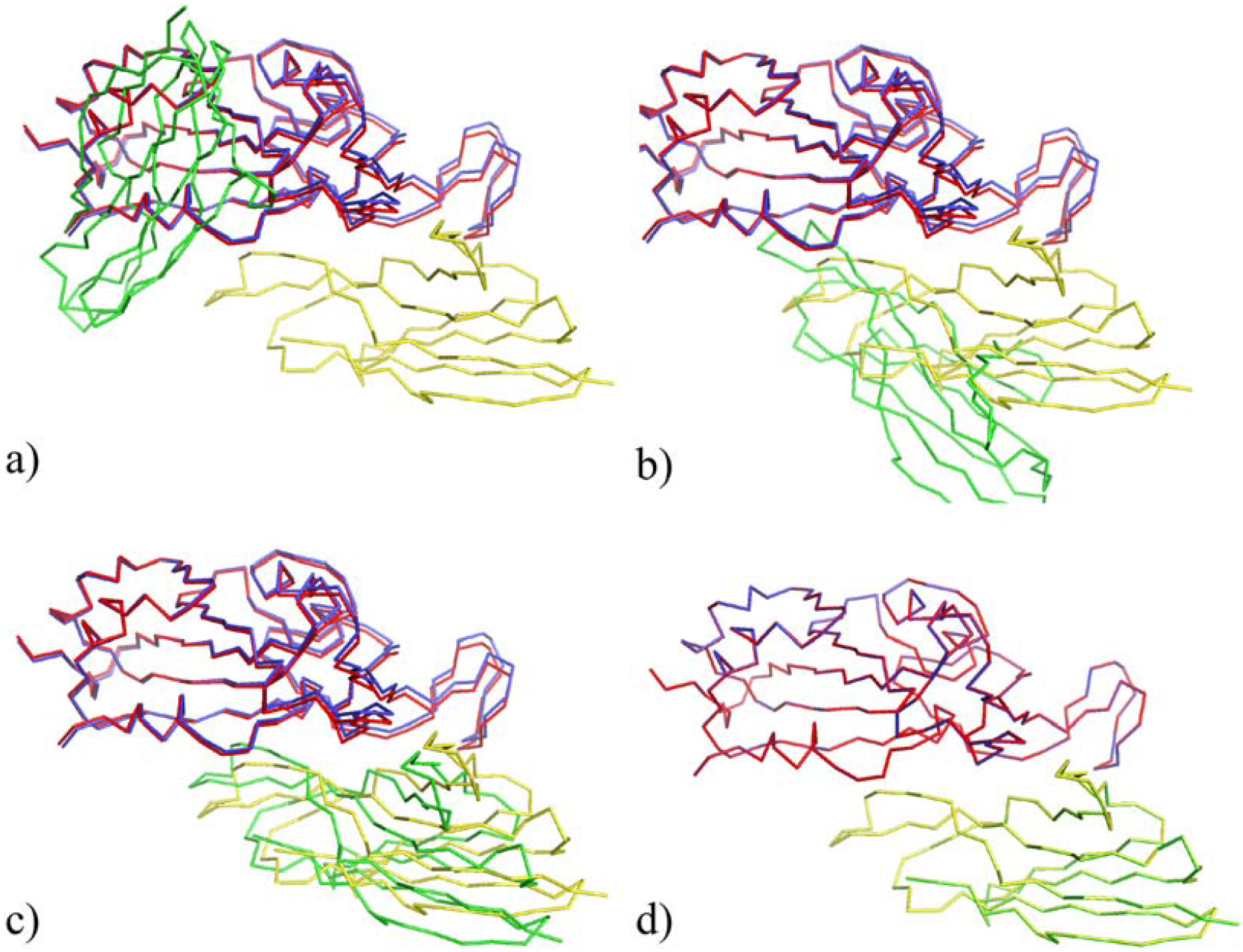
The intermediate structures of hetero-oligomer target 7SPP during the protein-protein docking process of OPUS-Fold3. The blue and yellow structures represent the native backbones of the receptor and ligand, respectively. The red and green structures correspond to their respective predicted intermediate structures. Images a)-d) illustrate the predicted intermediate structures at epochs 0, 40, 50, and 200, respectively.

Additionally, we evaluate the performance of OPUS-Fold3 on simultaneously modeling the backbones of the receptor and ligand, as well as the docking pose between them from scratch (OPUS-Fold3 (bbfold) in Table 5). Here, random values are set as the initial backbone torsion angles (*Φ* and *Ψ*). As an example, the trajectory of OPUS-Fold3 (bbfold) on hetero-oligomer target 7SPP is displayed as a movie in Supplementary Video S3. The results also suggest that the all-atom folding and docking procedure (OPUS-Fold3 (aafold) in Table 5) may achieve better performance than that using constraints from the backbone exclusively (OPUS-Fold3 (bbfold) in Table 5).

For further illustration, we present a folding and docking trajectory on hetero-oligomer target 7VNB as a movie in Supplementary Video S4. The backbones of the receptor and ligand are initially known, and random values are set as the initial side-chain dihedral angles (*X*_*1*,_ *X*_*2*,_ *X*_*3*,_ and *X*_*4*_). OPUS-Fold3 docks the receptor and ligand in the first 200 epochs, models the side chains in the following 2400 epochs, and finally performs folding and docking simultaneously on all atoms during the last 600 epochs as relaxation. The results indicate that when using a given set of constraints, OPUS-Fold3 can accurately deliver its corresponding 3D structure as a folding and docking framework.

## Concluding Discussion

In this paper, we introduce OPUS-Fold3, a gradient-based protein all-atom folding and docking framework. It is capable of integrating information from different sources such as physicochemical potentials, structure prediction techniques, cryo-EM data, and the constraints obtained by other methods. Our results show that, it can incorporate distance and orientation distribution constraints on the backbone folding task and achieve performance comparable to pyRosetta. Also, it can incorporate experimental cryo-EM maps as a set of constraints to improve the correlations between sidechains and densities in experimental maps.

In comparison to protein prediction methods like AlphaFold2 [1], which produce a specific structure based on multiple sequence alignment (MSA), folding framework like OPUS-Fold3 can be flexibly utilized to generate a protein 3D structure following multiple sources of constraints, which is crucial for protein structure refinement and protein design. Furthermore, compared to traditional schemes such as CNS [2] and pyRosetta [3, 4], OPUS-Fold3 is written in Python and TensorFlow 2.4, allowing it to be conveniently integrated with other deep learning models, bridging the gap between the traditional computational biology and AI communities.

## Acknowledgements

The work was supported by Shanghai Municipal Science and Technology Major Project (No.2018SHZDZX01), and ZJLab. Jianpeng Ma wants to thank the supports from National Key Research and Development Program of China (No. 2021YFF1200400), and Shanghai Artificial Intelligence Lab (P22KN00271). Ruhong Zhou wants to thank the support from the National Key R&D Program of China (2021YFA1201200 and 2021YFF1200404), the National Natural Science Foundation of China (U1967217), the National Independent Innovation Demonstration Zone Shanghai Zhangjiang Major Projects (ZJZX2020014), the National Center of Technology Innovation for Biopharmaceuticals (NCTIB2022HS02010), Shanghai Artificial Intelligence Lab (P22KN00272), and the Starry Night Science Fund of Zhejiang University Shanghai Institute for Advanced Study (SN-ZJU-SIAS-003).

## Conflict of Interest

None declared.

## Key Points

- We develop a gradient-based protein all-atom folding and docking framework, named OPUS-Fold3, which is able to accurately generate protein 3D structures in compliance with specified constraints, as long as the constraints can be expressed as functions of heavy atoms’ positions.
- In OPUS-Fold3, experimental cryo-EM density map can be included as a differentiable constraint and be integrated with other constraints such as those derived from structure prediction methods to jointly guide the optimization process, thus making a brige between the reconstruction of cryo-EM density map and protein structure prediction techniques.
- OPUS-Fold3 can be flexibly utilized to generate a protein 3D structure following multiple sources of constraints, which is crucial for protein structure refinement and protein design. Moreover, developed using Python and TensorFlow 2.4, OPUS-Fold3 is user-friendly for any source-code level modifications and can be seamlessly combined with other deep learning models, thus facilitating collaboration between the biology and AI communities.

## Notes

### Competing Interest Statement

The authors have declared no competing interest.

### Summary of Updates

1.We develop a gradient-based protein all-atom folding and docking framework, named OPUS-Fold3, which is able to accurately generate protein 3D structures in compliance with specified constraints, as long as the constraints can be expressed as functions of heavy atoms' positions. 2.In OPUS-Fold3, experimental cryo-EM density map can be included as a differentiable constraint and be integrated with other constraints such as those derived from structure prediction methods to jointly guide the optimization process, thus making a brige between the reconstruction of cryo-EM density map and protein structure prediction techniques. 3.OPUS-Fold3 can be flexibly utilized to generate a protein 3D structure following multiple sources of constraints, which is crucial for protein structure refinement and protein design. Moreover, developed using Python and TensorFlow 2.4, OPUS-Fold3 is user-friendly for any source-code level modifications and can be seamlessly combined with other deep learning models, thus facilitating collaboration between the biology and AI communities.

